# Epistasis and the dynamics of reversion in molecular evolution

**DOI:** 10.1101/042895

**Authors:** David M. McCandlish, Premal Shah, Joshua B. Plotkin

**Affiliations:** Department of Biology, University of Pennsylvania, Philadelphia, PA; Department of Genetics, Rutgers University, Piscataway, NJ

**Keywords:** weak mutation, fitness landscape, entrenchment, reversible Markov chain

## Abstract

Recent studies of protein evolution contend that the longer an amino acid substitution is present at a site, the less likely it is to revert to the amino acid previously occupying that site. Here we study this phenomenon of decreasing reversion rates rigorously, and in a much more general context. We show that, under weak mutation and for arbitrary fitness landscapes, reversion rates decrease with time for any site that is involved in at least one epistatic interaction. Specifically, we prove that, at stationarity, the hazard function of the distribution of waiting times until reversion is strictly decreasing for any such site. Thus, in the presence of epistasis, the longer a particular character has been absent from a site, the less likely the site will revert to its prior state. We also explore several examples of this general result, which share a common pattern whereby the probability of having reverted increases rapidly at short times to some substantial value before becoming almost flat after a few substitutions at other sites. This pattern indicates a characteristic tendency for reversion to occur either almost immediately after the initial substitution or only after a very long time.

## Introduction

In the context of evolutionary theory, reversion describes a population that returns to an ancestral character state (Porter and Crandall 2003). While many early (Dollo 1893; Muller 1939; Simpson 1953; Gould 1970) and more recent (Teotonio and Rose 2001; Collin and Miglietta 2008; Bridgham et al. 2009; Tan et al. 2011) discussions of reversion consider an environmental change that confers a selective advantage to an ancestral phenotype, reversion may also occur at the level of nucleic acids or protein sequences, with evolution proceeding under longterm purifying selection (Kimura 1983). Such reversions occur both because of the strictly limited number of character states (four possible nucleotides or twenty possible amino acids, Jukes and Cantor 1969) and because selection on molecular function may constrain a given position to only a subset of these possible character states (Rokas and Carroll 2008; Breen et al. 2012).

It has long been hypothesized that epistatic interactions should lower the rate of reversion, rendering evolution effectively irreversible (Muller 1918, 1939). This issue has been especially important recently, due to ongoing debate in the field of protein evolution about how position-specific preferences for amino acids may change over time (Pollock et al. 2012; Naumenko et al. 2012; Ashenberg et al. 2013; Pollock and Goldstein 2014; Risso et al. 2015; Shah et al. 2015; Usmanova et al. 2015; Goldstein et al. 2015; Bazykin 2015). In particular, several groups have suggested that once an amino acid substitution occurs at a particular position, epistatic interactions with subsequent substitutions at other positions should tend to increase the selective preference for the derived amino acid relative to the ancestral state (Pollock et al. 2012; Naumenko et al. 2012; Shah et al. 2015, c.f. Fisher 1930, pg. 95), a phenomenon known as entrenchment (Shah et al. 2015). This means that a mutation that was nearly neutral when it originally went to fixation may become increasingly deleterious to revert, which would cause a decreasing propensity to revert as time elapses.

However, the above verbal argument is not entirely convincing. While it is easy to imagine some forms of epistasis that would cause reversion rates to decrease over time, evolutionary dynamics on high-dimensional fitness landscapes can have many counter-intuitive properties (Conrad 1990; Gavrilets 1997; Carneiro and Hartl 2010; Kondrashov and Kondrashov 2015; McCandlish et al. 2013, 2015b). Here we undertake a rigorous mathematical investigation into the relationship between the rate of reversion and the presence of epistasis, for arbitrary fitness landscapes. We study this problem under the assumption that mutation is weak relative to drift, so that the evolution of the population can be modeled as a Markov chain on the set of genotypes (for a review see McCandlish and Stoltzfus 2014). Our main conclusion is that, for any site involved in at least one epistatic interaction, the rate of reversion for a substitution is a strictly decreasing function of the time since the initial substitution.

Our first task is to provide a rigorous definition for the rate of reversion. For a population evolving under weak mutation, we can consider the population as a single particle that jumps from one genotype on the fitness landscape to another at each substitution event. We consider some focal set of genotypes that includes the starting state of the population. If we observe the population for long enough, the population will eventually leave this focal set and trace a path through the space of genotypes. At each point along this path, it has some propensity to fix a genotype in the focal set, that is, to revert. Eventually, this propensity is realized and the population returns to the focal set. If we continue to watch the population for long enough, this process will repeat itself many times, and we can ask the question: given that the population left the focal set t time units ago and has not yet returned to it, what is the instantaneous expected propensity for that population to return to the focal set, i.e. what is the rate of reversion as a function of time?

To study reversion it is helpful to note the following relationship between the rate of reversion and the distribution of waiting times until a reversion event occurs. As we observe the population evolve on the fitness landscape, every time the population leaves the focal subset, we can record the waiting time until it first returns. And again, if we observe the population for long enough, these waiting times will converge to a particular distribution. Importantly, the reversion rate described above is equal to the hazard function of this probability distribution of reversion times, that is, the probability density of this distribution at time t, conditioned on drawing a value t or greater. Thus, we can study how the rate of reversion changes the longer the population has been absent from the focal subset by studying the hazard function of this distribution of reversion times.

The case of a bi-allelic non-epistatic (i.e. additive) fitness landscape provides an instructive, introductory example. In this case, it is easy to show that the distribution of return times for a particular allele at a particular site is always exponentially distributed, corresponding to a constant hazard function. That is, for a bi-allelic site on a non-epistatic fitness landscape, the reversion rate does not change as a function of time. We want to understand how this simple situation changes in the presence of epistasis.

Here we show that if the focal site interacts epistatically with at least one other site, then the hazard function of the distribution of reversion times, and therefore the rate of reversion, is strictly decreasing in time. This implies that the longer a population has been away from the focal set of genotypes, the longer the expected waiting time until it returns to that set. Moreover, this decreasing reversion rate is due to two factors, each of which would individually result in a decreasing reversion rate.

The first factor is co-evolution between sites as suggested by, e.g., Pollock et al. (2012). As long as the derived allele is resident at the focal site, it forms part of the genetic background for other substitutions, and this causes the population to tend to spend more time at genotypes where the derived allele is selectively favored.

To isolate the effect of this first factor, we consider a modified process where we do not allow the focal site to return to its original state after the initial substitution. Thus, the dynamics after the initial substitution capture the acclimatization of the rest of the genome to the derived state at the focal site. While reversion events cannot occur under this modified model, we nonetheless keep track of the reversion rate that would occur if we were to suddenly allow reversions. We show that the reversion rate for this modified process is decreasing for sites involved in at least one epistatic interaction, which demonstrates that co-evolution between sites leads to reversion rates that decrease in time.

The second factor that produces decreasing rates of reversion is statistical in nature. This second factor arises because in order to focus on the first time a population returns to the ancestral state we must condition on that return not having yet occurred when we calculate the rate of reversion. If the population is at a genotype with a high reversion rate, it tends to actually revert, so that the high reversion rate no longer contributes to the expectation. This alone results in reversion rates that decrease in time. To put this in a more biological light, populations that have been gone for a long time from the focal subset tend to have low propensity to return to the focal subset, because if they had a high propensity they would have returned already.

To isolate the effects of this second factor, we can consider a different, modified model in which the population never moves to another genotype outside the focal set once it leaves the focal set. Thus, each time the population leaves the focal set, its propensity to return to the focal set is constant. Nonetheless, the reversion rate under this model will be strictly decreasing in time if there is any variation among these genotypes in the propensity to return to the focal set.

In actuality, these two factors operate simultaneously, and their effects on the time-evolution of the rate of reversion are coupled. Nonetheless, when both factors operate, we will show that the reversion rate is still decreasing with the time since the substitution of the derived allele at the focal site.

We will also consider what occurs when more than two alleles are available at a site and the more general case of reversion to subsets of genotypes, e.g. reversion to the set of codons corresponding to a particular amino acid. The key observation in this context is that the two factors above operate when there is any genotype-to-genotype variation in the propensity to return to the focal set. While for models with bi-allelic sites epistasis is the only way of producing variation in these propensities, for more general models with more than two alleles per site the rate of reversion may be decreasing even in the absence of epistasis.

In addition to our main results, which concern populations that have already been evolving on the same fitness landscape for a long time, we will briefly explore how changes to the fitness landscape effect the dynamics of reversion. Finally, we will explore several simple examples to gain intuition for the magnitude and evolutionary importance of reversion rates that decrease in time.

## Materials and methods

### Population-genetic model

We consider a population evolving in continuous time under weak mutation on an arbitrary finite-state fitness landscape (e.g. Iwasa 1988; Sella and Hirsh 2005; McCandlish et al. 2015b). In this regime, we can model the population as a single particle that moves from genotype to genotype at each fixation event (see McCandlish and Stoltzfus 2014, for a review). More formally, we model evolution as a continuous time Markov chain with a rate matrix **Q**_full_:

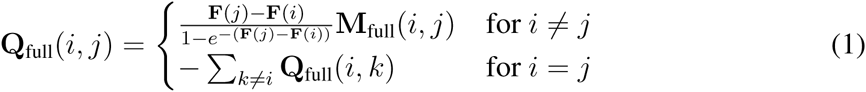

where **F**(*i*) is the fitness of genotype *i* and **M**_full_(*i*, *j*) is the mutation rate from *i* to *j*. We further assume that the fitness landscapes is connected, so that there exists a mutational path between any two genotypes *i* and *j* and that the Markov chain defined by **Q**_full_ is reversible, so that there exists a stationary probability distribution **π**_full_ of the chain defined by **Q**_full_ such that ***π***_full_(*i*)**Q**_full_(*i*, *j*) = ***π***_full_(*j*)**Q**_full_(*j*, *i*) for all *i*, *j*. This latter condition will be satisfied whenever the neutral mutational dynamics produce a reversible Markov chain (see e.g., Sella and Hirsh 2005; McCandlish et al. 2015b); a simple sufficient condition is that the mutation rates are pairwise symmetric, **M**(*i*, *j*) = **M**(*j*, *i*) for all *i*, *j*.

We are interested in the situation where a population has just left some subset of states *A* and want to study the waiting time for the population to return to that subset *A*. Without loss of generality, we can order the states so that all states in *A* come after the states in the complement of *A*, so that we can write **Q**_full_ in block matrix form as:

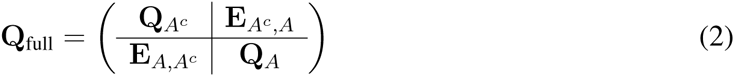

where *A*^*c*^ is the complement of *A* and **E**_*A*^*c*^,*A*_ gives the transition rates from *A*^*c*^ to *A* and **E**_*A*,*A*^*c*^_ gives the transition rates from *A* to *A*^*c*^.

Our main object of study will be the absorbing Markov chain with rate matrix **Q**_*A*^*c*^_, where absorption corresponds to a return to the subset *A*. Because we will assume that a population starts at time 0 having just left the subset of states *A*, this means that an absorption event is also a reversion event, so that we can study the dynamics of reversion by studying the waiting time until absorption for the Markov chain with rate matrix **Q**_*A*^*c*^_. For brevity, we will simply call this matrix **Q**.

The row sums of –**Q** (or equivalently, the row sums of **E**_*A*^*c*^,*A*_) give the propensity for a population currently fixed at genotype *i* to return to *A*, and we write the rate at which such an event occurs for a population fixed at genotype *i* as **g**(*i*). Let x_*t*_(*i*) be the probability that the population is fixed for genotype *i* and has not yet reverted at time *t*. Then the time evolution of *x*_*t*_ is given by

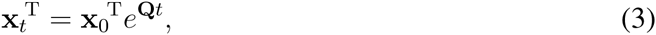

where x_0_(*i*) gives the probability that the population initially left the subset *A* by becoming fixed for genotype *i*.

### The hazard function of reversion times

Consider a population that first leaves subset *A* by fixing genotype *i*. The probability that the population reverts, that is first becomes fixed for a genotype in the subset *A*, during the time interval [*t*, *t* + *dt*) is given by *f*_*i*_(*t*) *dt* where

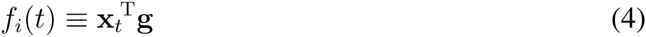

and *x*_*0*_(*j*) is 1 for *i* = *j* and 0 otherwise. Thus, *f*_*i*_(*t*) is the probability density function of the distribution of reversion times for a population that initially leaves subset *A* by fixing genotype i; we note that *f*_*i*_(*t*) is indeed a proper probability density since **Q**_full_ defines an ergodic Markov chain and so populations return to the subset *A* with probability 1.

Now, a population that has already been evolving on a fitness landscape for a long time is much more likely to leave the subset *A* by fixing some genotypes rather than others. To capture this effect, we can choose the initial distribution x_0_ by considering a population whose genotype is described by the stationary distribution ***π***_full_ and then condition on leaving the subset *A* in the interval [0, *dt*). We thus specify the distribution x_0_ as

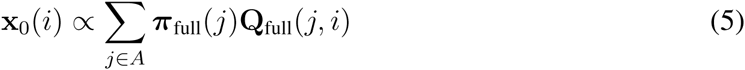

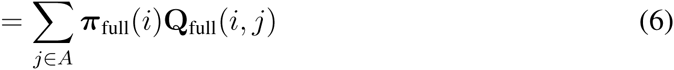

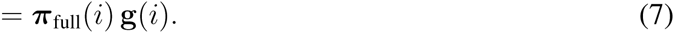

With this choice of initial distribution, the probability density function for the distribution of reversion times is given by

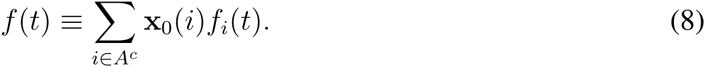

Note that this is the same distribution that we would get if we watched a single population evolve for an infinite amount of time and recorded, each time the population left *A*, the waiting time to return to *A*.

We now turn to formally defining the rate of reversion. What we want to understand is how the rate at which a population first returns to some set of states changes the longer a population has been outside that set. This suggests that we should define the reversion rate as the probability density of a population returning to set *A* for the first time in the time interval [*t*, *t* + *dt*) given that the population has not already returned to set *A* before time *t*. In the more general context of non-negative probability distributions, this quantity is known as the hazard function (or failure rate, or force of mortality). Thus, we define the reversion rate to be hazard function of the probability distribution of reversion times.

For instance, consider the distribution of reversion times at stationarity, with density given by *f*(*t*), cumulative distribution function 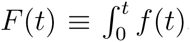, and complementary cumulative distribution function *F̄*(*t*) ⁡ 1 – *F*(*t*). Then the reversion rate at time *t* is given by the hazard function *h*(*t*) where

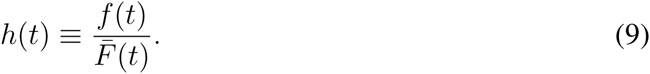

If *h*(*t*) is increasing in *t* it means that populations that have been away from the set *A* for a long time on average have instantaneous rates of return to a larger than the average rate of return to *A* of populations that have only been away for a short time. Likewise, if *h*(*t*) is decreasing in t it means that populations that have been away from the set *A* for a long time on average have instantaneous rates of return to *A* smaller than the average rate of return to *A* of populations that have only been away for a short time. Another quantity of interest is the expected remaining time until reversion, given that the population has not yet reverted at time *t*. This expected waiting time can be expressed in terms of the hazard function as

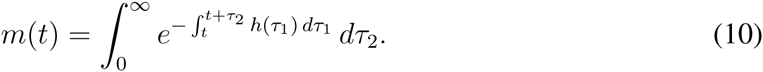

Furthermore, from the above equation it is easy to see that if the reversion rate is increasing in time, then the expected remaining waiting time is decreasing in time, whereas if the reversion rate is decreasing in time, then the expected remaining waiting time is increasing.

## Results

### General theory of reversions

Our first main result is that for a population that has already been evolving on the same fitness landscape for a long time, the rate of reversion to *A*, *h*(*t*), is a non-increasing function of t, where t is the time since the population left *A*. Moreover, the reversion rate *h*(*t*) is strictly decreasing unless all genotypes is *A*^*c*^ have the same reversion rate (i.e. **g**(*i*) is constant) in which case *h*(*t*) is also constant. This result shows that populations that have spent a longer time away from *A* cannot possibly have higher rates of return to the ancestral character state or shorter expected remaining waiting times until return.

This result is a simple consequence of the fact, well-known in the mathematical literature (Kielson 1979; Aldous and Fill 2002), that the distribution of return times to a subset for a stationary, finite-state, reversible, continuous-time Markov chains take the form of a mixture of exponential distributions (in the mathematical literature, such a distribution is known as a “completely monotone” distribution). It is easy to show that the hazard function for such a distribution is strictly decreasing except in the case of a pure exponential distribution. In Appendix 1, we provide elementary proofs of these facts as well as the fact that that the distribution of return times is a pure exponential only in the case when **g**(*i*) is constant.

### Connection to epistasis

An immediate implication of this result is that the reversion rate for any site in a fitness landscape that is involved in an epistatic interaction must be strictly decreasing in time. This is because epistatic interactions result in variation in the genotype-specific rates of return to *A*, **g**(*i*), since different back-mutations have different fitness consequences.

More formally, consider a bi-allelic fitness landscape with *L* sites and with forward and backward mutation rates of *μ*_*i*_ and *v*_*l*_, respectively, at site *l*. Thus, **M**_full_(*i*, *j*) = *μ*_*l*_ if *j* differs from *i* a forward mutation at site *l*, **M**_full_(*i*, *j*) = *ν*_*l*_ if *j* differs from *i* by a back mutation at site *l* and **M**_full_(*i*, *j*) = 0 otherwise. Furthermore, let us pick a focal site *l*\* and order the states such that all genotypes that have the allele produced by the forward mutation are in set *A* and indexed first. We are interested in the distribution of reversion times at site *l*\*.

We say a site is not involved in any epistatic interactions if its fitness effect is constant, in the sense that if genotype *i* differs from genotype *j* by a forward substitution then the scaled selection coefficient **F**(*i*) – *F*(*j*) is equal to some constant *S*_*l*_*. In this case **g**(*i*) is constant and equal to *v*_*l*\*_(*i*)*S*_*l*\*_/(1 – *e*^‒*S*_*l*\*_^), so that *h*(*t*) is constant. On the other hand, suppose *l*\* is involved in an epistatic interaction. Then there exist *i*, *i*′ ∈ *A*^*c*^ and *j*, *j*′ ∈ *A* such that both *i* differs from *j* and *i*′ differs from *j*′ by a forward mutation at site *l*\* but **F**(*i*) – **F**(*j*) = **F**(*i*′) – **F**(*j*′). But then **g**(*i*) = **g**(*i*′) because **Q**_full_(*i*, *j*) = *v*_*l*\*_(**F**(*j*) – **F**(*i*))/(1 – *e*^‒(**F**(*j*^)**F**«)) ≠ *v*_*l*\*_.(**F**(*j*′) – **F**(*i*′))/(1 – *e*^‒(**F**(*j*/)–**F**(*i*/))^) ≠ **Q**_full_(*i*′, *j*′). Thus, in this case *h*(*t*) is strictly decreasing.

The above argument shows that *h*(*t*) is strictly decreasing for a biallelic fitness landscape if and only if the focal site is involved in at least one epistatic interaction. For fitness landscapes that include more than two alleles at a site, by contrast, the notation and argument become somewhat more involved (Appendix 2), but the end result is weakened to the statement that epistasis is sufficient for *h*(*t*) to be strictly decreasing. Indeed, even for non-epistatic multiallelic fitness landscapes we generically expect *h*(*t*) to be strictly decreasing. This is because even if fitness is additive between sites, the different alleles within a site will have different fitness differences from the focal allele and therefore different probabilities of fixation that result in a non-constant **g**(*i*).

### Do reversions also become more deleterious?

Besides the relationship between decreasing reversion rates and epistasis, it is interesting to ask about the relationship between decreasing reversion rates and the mean selection coefficient of reversions. This relationship is not necessarily simple. First, this is because different genotypes might have different mutation rates to the subset *A*, so that the decrease in *h*(*t*) might be realized by the non-reverting subset of the population becoming concentrated at genotypes with low mutation rates to the subset *A* rather than by having more negative selection coefficients for such mutations. However, even if all genotypes in *A*^*c*^ produce mutations to subset *A* at the same rate the decrease in *h*(*t*) might not correspond to reversion mutations becoming more deleterious. This is due to the non-linearity of the function relating the scaled selection coefficient S to the rate of evolution; in particular if *S* is the scaled selection coefficients, this function is *K*(*S*) = *S*/(1 – *e*^‒*S*^). However, using the fact that *K*(*S*) is concave (McCandlish et al. 2015a), if each genotype in *A*^*c*^ produces mutations at rate *μ* to a single corresponding genotype in set *A*, then Jensen’s inequality tells us that *h*(*t*) is greater than *μ* times *K*(*S*_*t*_), where *S*_*t*_ is the average scaled selection coefficient of the reversion at time t and this average is taken with respect x_*t*_/*F*(*t*). Thus, if for any t we have *h*(*t*) < *μ*π*K*(*S*_0_), then the mean selection coefficient of a reversion among those populations that have not yet reverted is less than or equal to the mean selection coefficient of a reversion immediately after the initial substitution away from *A*. This provides a sufficient condition for the mean selection coefficient of reversions to have decreased in time.

### Expected reversion times

If the reversion rate is non-increasing over time, then the expected waiting time until reversion must be greater than what would be predicted based on the initial reversion rate alone, i.e. we must have *m*(0) ≥ 1/*h*(0). In fact, there is a simple formula for the expected time until reversion that holds even if the Markov chain describing the weak mutation dynamics is not reversible. In particular, consider the expected rate of returns to *A* at stationarity when we condition on a population being in the subset *A*^*c*^, i.e. ∑_*i*∈*A*_*c*__ ***π***_full_(*i*)**g**(*i*)/∑_*i*∈*A*_*c*__ ***π***_full_(*i*). Then *m*(0) is simply the reciprocal of this rate (see Appendix 3).

However, because the reversion rate is non-increasing in time, this mean value does not necessarily provide a very informative summary of the dynamics of reversion. For instance, consider the case where *h*(*t*) is initially high, but drops to a very low value. Under these circumstances, it can be the case that a large proportion of populations revert rapidly, but the subset of populations that do not revert rapidly may have extremely long expected reversion times, so that the mean reversion time *m*(0) is really an average over two very different subsets of populations. Intuitively, this situation will often arise when the set a is the basin of attraction of a fitness peak. Most populations that cross into the fitness valley around a will return to *A* rather than cross the valley, but a small subset will cross the fitness valley and end up at another fitness peak. The expected return time for this subset of populations might then be extremely long.

### Why is the reversion rate decreasing?

In order to gain an intuitive understanding for why the reversion rate is decreasing in time, it is helpful to distinguish between two different phenomena that each contribute to this decrease.

The first phenomenon is the acclimatization of the rest of the genotype to being in the subset *A*^*c*^. For instance, in the context of protein evolution, once a mutation has fixed at a focal site it forms part of the genetic background that determines the fitness effects of mutations at other sites. For sites that interact epistatically with the focal site, this will tend to favor substitutions whose effects are more positive when the derived allele is present at the focal site.

Such acclimatization of the genome tends to result in a decreasing reversion rate at the focal site. To make this idea precise, let us consider a modified process where we do not allow populations to revert to *A* after they enter the subset *A*^*c*^. Following this initial entrance, the dynamics are thus governed by a rate matrix **Q**\* where **Q**\* ≡ **Q** + **D**_g_ and **D**_g_ is the diagonal matrix whose *i*, *i*-th entry is *g*(*i*). For ease of exposition, let us also assume that the subset *A*^*c*^ is connected (i.e. one can always evolve from any state in *A*^*c*^ to any other state in *A*^*c*^ without returning to *A*; the more general case is handled in Appendix 1).

Even though we do not allow reversions to occur back to a under this modified process, we can still keep track of the reversion rate that would occur were we to allow reversions. When the population initially leaves the subset *A*, it is distributed as x_0_(*i*) ∝ ***π***_full_(*i*)**g**(*i*). However, as time elapses under the modified process, the probability that the population is fixed for genotype *i* ∈ *A*^*c*^ tends to a distribution that is ∝ π_full_(*i*). Now, the initial reversion rate under the modified process is equal to the expected value of **g**(*i*) with respect to the first of these distributions, while the asymptotic reversion rate under the modified process is equal to the expected value of **g**(*i*) with respect to the second. Clearly, the reversion rate is higher under the first distribution, since this distribution differs from the second only in that it is more concentrated at values with high **g**(*i*). Indeed in Appendix 1 we show the stronger result that the reversion rate is strictly decreasing for the modified process unless **g**(*i*) is constant, in which case the reversion rate is also constant.

The second phenomenon is a statistical phenomenon having to do with the fact that we are interested in the first return to *A*. Even if there was no acclimatization to the initial substitution (and therefore no co-evolution) the reversion rate would tend to be decreasing in time due to genotype-specific variation in the rates of return to a, i.e. the **g**(*i*). This effect occurs because populations that have not reverted even after a long time have likely spent most of this time at genotypes where returns to *A* are unlikely. To put this another way, if the population had spent a great deal of time at genotypes with high return rates, then it would have returned already. This phenomenon is well-known in the reliability and demography literature, where unaccounted for heterogeneity can result in failure or mortality rates that decrease in time even when individual failure or mortality rates are constant (Vaupel and Yashin 1985).

To isolate this second phenomenon, we consider a second modified process in which a population that leaves *A* by fixing genotype *i* experiences no additional substitutions until it returns to *A*, i.e. the return rate for such a population is always **g**(*i*). The rate matrix for this process is thus –**D**_g_. The derivative of the reversion rate of this process is equal to –1 times the variance in g conditional on not having yet reverted. Since variances are non-negative, the derivative of the reversion rate is non-positive, so that the reversion rate is non-increasing.

While we have constructed these two modified processes to separate the effects of acclimatization of the genome and the effects of conditioning on not having yet reverted, these two effects interact to determine the dynamics of the original process. The fact that both phenomena tend to lead to decreasing reversion rates when there is variation in g (or more precisely for the second phenomenon, variation in its non-zero elements), helps clarify why the reversion rate is non-increasing for the original process.

### Reversion rates under non-stationary evolution

So far, we have concentrated on the case of a population that has already been evolving on a fixed fitness landscape for a long time, so that each possible way of leaving the subset *A* is weighted by its stationary probability. However, we can also gain some strong intuitions for the behavior of the reversion rate in the more general case, where we allow the initial probability vector x_0_ to be arbitrary.

In particular, in this more general setting we can show that the time-evolution of the (sign-reversed) reversion rate is isomorphic to the time-evolution of the mean fitness of an infinite population on a suitable fitness landscape. The basic idea is that the time-evolution of the probability distribution describing the genotype of a population conditional on not having yet reverted can be viewed as the time-evolution of the frequencies of genotypes in an infinite population whose mutational dynamics are specified by the off-diagonal entries of the matrix Q and where the Malthusian fitness of genotype *i* is given by –**g**(*i*). That is, **g**(*i*) plays the role of a genotype-specific death rate.

More formally, let p_*t*_(*i*) be the probability that a population is fixed for genotype *i* at time *t* given that it has not returned to the subset *A* by time *t*. Then p_*t*_(*i*) = x_*t*_(*i*)/∑_*j*_x_*t*_(*j*). Writing *i* for the identity matrix𝟙 for the vector of all 1s, and **D**_‒g_ for the diagonal matrix with ‒g down its main diagonal, we have

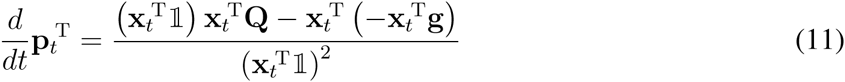

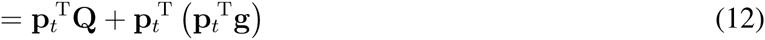

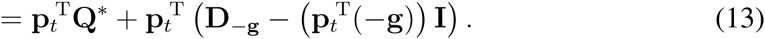

This is simply the “parallel” version of the replicator equation (i.e. where mutation occurs independently from reproduction), with **Q**\* as the rate matrix of the mutational process and ‒g as the vector of fitnesses. Furthermore, the reversion rate at time *t* is then given by p_*t*_^T^g, which is simply the mean fitness p_*t*_^T^(–g) with its sign reversed. The derivate of the reversion rate is then:

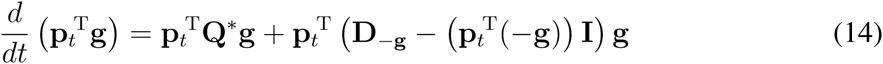

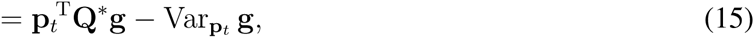

where Var_**p**t_ g is the variance in the return rate with respect to the probability distribution p_*t*_. Here, the term p_*t*_^T^**Q**\*g gives the effect on the reversion rate due to acclimatization of the genotype to being in the subset *A*^*c*^, while – Var_pt_ g captures the effects of conditioning on not having yet reverted. Unlike under stationarity, the effects of acclimatization for the non-stationary case can either increase or decrease the reversion rate. However, conditioning on not having yet reverted still always produces a bias towards a decreasing reversion rate, since ‒Var_pt_ g is non-positive.

## Examples

The preceding results describe a general tendency for the reversion rate–quantified as the hazard function of the distribution of return times–to be decreasing in the time. Here we present some simple examples to explore the magnitude and evolutionary consequences of this effect.

### Crossing fitness valleys

Consider the case of a biallelic fitness landscape with a symmetric fitness valley. In particular, consider the case where genotypes ab and AB have equal fitnesses and scaled selection coefficient S with respect to the valley genotypes Ab and aB and mutations occur independently at each locus at rate 1. Let us now examine the rate of reversion at the first site, so that the set *A* = {ab, aB} and leaving *A* corresponds to an a→A substitution. Given that the population has already been evolving on this fitness landscape for a long time and that an a→A substitution has just occurred, we want to understand the distribution of times until an A→a substitution occurs as a function of the depth of the fitness valley *S*.

This distribution of times is easy to understand intuitively. For S less than ≈ 2 the dynamics are approximately neutral and the waiting time for reversion is approximately exponential with mean 1. For larger *S*, the dynamics are substantially affected by the fitness valley. The best way to understand these dynamics is by considering where the population is immediately after an a→A substitution. At stationarity, we know that the frequency of substitutions into the fitness valley must be equal to the frequency of substitutions out of the valley. This means that at stationarity, half the time the population will have just fixed the valley genotype *Ab* and half the time it will have fixed the peak genotype *AB*. Populations that fix the peak genotype AB are unlikely to revert in the short-term, whereas populations that fix the valley genotype Ab are likely to move to one or the other peak in the short-term, with equal probability. Thus, roughly speaking, after a short time the population will either have reverted (with probability 1/4, since half the time it fixes the valley genotype and then immediately reverts half the time) or be fixed at the fitness peak AB (with probability 3/4). This means that reversions happen either very shortly after the initial substitution—and this occurs 1/4 of the time—or only after the long waiting time needed for deleterious fixations to occur.

Figure 1 shows these dynamics as a function of *S*. The top row of panels shows the probability that an A→a reversion has occurred as a function of time and the bottom row of panels shows the reversion rate, *h*(*t*). The left-most column show the neutral case. The reversion rate is constant and the probability that a reversion has occurred by time t is given by 1 – *e*^‒*t*^. The middle and right columns show the case where *S* = 2.5 and *S* = 5 respectively. In both cases, the reversion rate is high initially, starting at approximately *S*/2 (probability 1/2 of starting at the valley genotype in which case returns to *A* occur at roughly rate *S*). The reversion rate drops rapidly as populations leave the valley genotype, approaching an asymptotic rate of approximately 3/2 the substitution rate of a deleterious fixation – *S*/(1 – e^S^) (populations that have not yet reverted are likely at the AB fitness peak; reversion occurs either via a direct substitution of a at rate – *S*/(1 – *e*^*S*^) or via fixation of the valley genotype Ab, which occurs at rate –*S*/(1 – *e*^*S*^), followed by reversion with probability 1/2). Thus, the reversion rate is initially high but rapidly approaches a much lower rate, which results in a characteristic “knee” in the probability that a population has reverted by time *t*, where a population has a substantial probability of having reverted at short times (corresponding to a steep initial increase in the fraction reverted as a function of time), but populations that do not revert during this initial period have much lower reversion rates (resulting in a slow increase in the fraction reverted after this initial perioed), see Figure 1 first row, middle and right columns.

**Figure 1:**
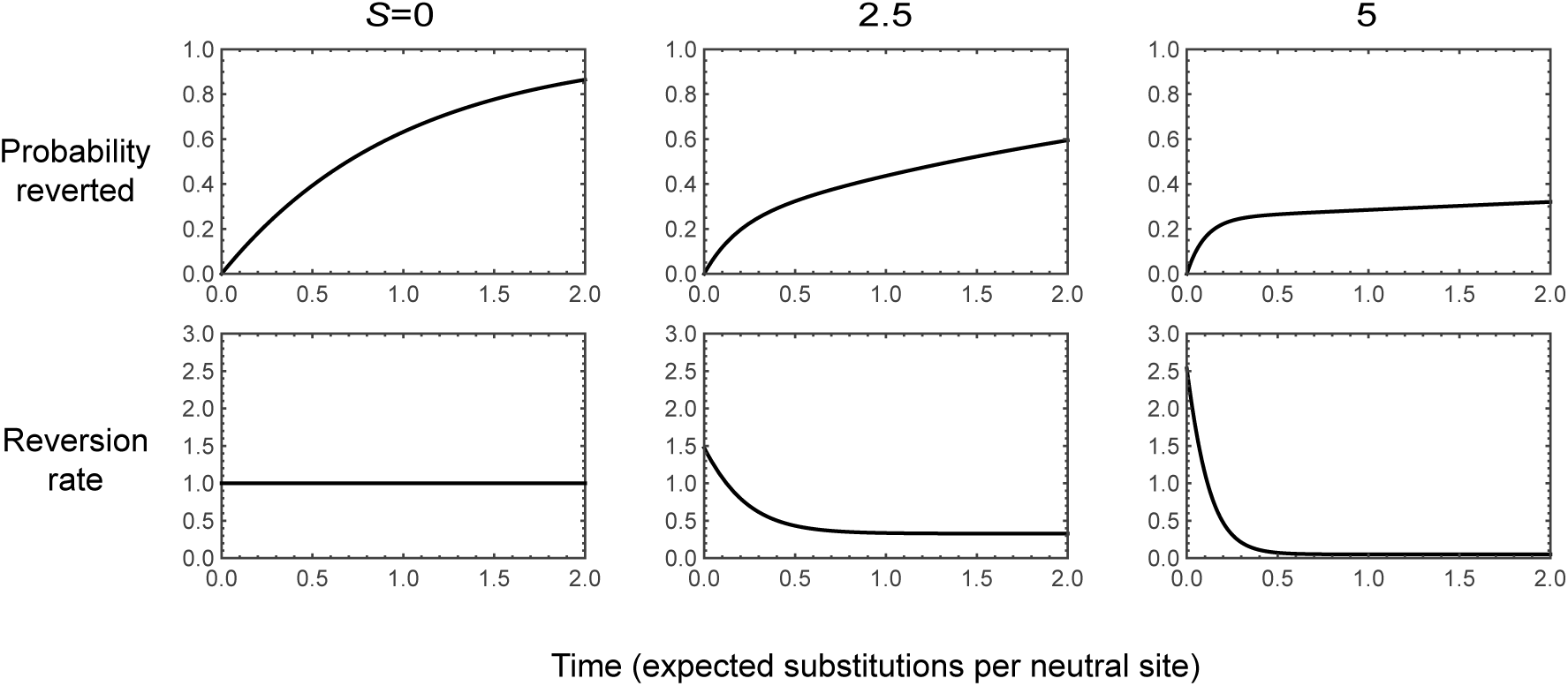
Reversion rates for a biallelic fitness landscape with a symmetric fitness valley. Genotypes ab and AB have equal fitness and scaled selective advantage *S* relative to genotypes Ab and aB, and mutations occur at each site at rate 1 in each direction. The dynamics are shown for returns to the focal subset *A* = {ab, aB} assuming that the population has just left the subset a at stationarity. Each column corresponds to a different value of S. The first row shows the probability that the population has reverted as a function of time, and the second row shows the reversion rate, *h*(*t*).

### Mutational robustness and the rate of reversion

As a second example, we consider a biallelic fitness landscape with one focal site and L other sites, where each genotype is viable with probability p and all viable genotypes are neutral relative to each other. All sites experience forward and reverse mutations at rate 1. This example is motivated by the model used by Kondrashov and colleagues to study long-term purifying selection (Breen et al. 2012; Usmanova et al. 2015). We want to understand the dynamics of reversion at the focal site.

We proceed with a heuristic treatment based on the assumption that *Lp* ≫ 1. In this regime, we treat each genotype as having a constant number *Lp* of neutral neighbors accessible by mutations at non-focal sites. The key idea is that immediately after a substitution at the focal site, the back-substitution is viable and so reversions initially occur at rate 1. However, once an additional substitution at a non-focal site accrues, the probability that the reversion mutation at the focal site is viable is only *p*. Thus, the reversion rate drops very rapidly from 1 to approximately *p*.

In particular, after a substitution at the focal site, the expected substitution rate is 1 + *Lp* and the probability that the first substitution is a reversion is 1/(1 + *Lp*). Thus, we can approximate the probability distribution of waiting times until reversion as a mixture of two exponential distributions. The first distribution has rate 1 and is weighted by the probability that revert before any other substitution occurs, 1/(1 + *Lp*). The second exponential distribution has rate p and is weighted by the probability that a population experiences a substitution at another site before the reversion event occurs, *Lp*/(1 + *Lp*). This suggests that the probability of having reverted by time t can be approximated by:

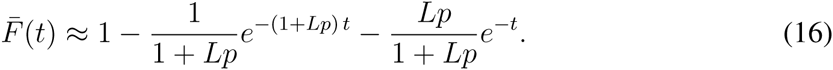

Based on a similar argument, we can approximate the reversion rate as

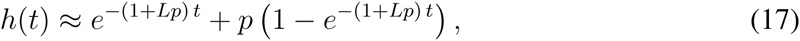

where the first term corresponds to the initial reversion rate of 1, which applies with probability approximately e^-(1+Lp)^ ^t^ (the probability of not having yet having experienced the first substitution) and the second term corresponds to the expected reversion rate given that one substitution has occurred, p, weighted by the probability that a substitution has already occurred, 1 *e*^‒(1+*Lp*)*t*^

Although an exact treatment of any particular realization of the model is possible using the associated rate matrix **Q**_full_, the construction of this matrix is not computationally feasible for large *L*. We thus proceeded by simulation. A population that has just experienced a substitution at the focal site must be at a viable genotype and have just come from a viable genotype that differs only at the focal site. We simulated the subsequent evolution of such a population under weak mutation, keeping track of the genotypes produced by mutation and whether they are viable (with probability *p*) or inviable (with probability 1 – *p*) until the first reversion event at the focal site. For each choice of parameters, we repeated this procedure 1,000,000 times.

The resulting reversion times are shown in Figure 2 for *L* = 100. The first row shows the probability of having reverted by time *t*, and the second row shows the reversion rate as a function of time. The first column shows the exact theoretical results for the case where all genotypes are viable (gray lines). The second and third columns show the results for *p* = .5 and *p* = .1, respectively, with the approximate theoretical results given by Equations 16 and 17 in gray and the simulation results in black. For the case *p* = .5 the reversion rate drops rapidly from 1 to approximately *p*. Indeed, this drop is so rapid that relatively few populations actually revert at the higher initial rate, which is as we expect, since the probability that a population reverts before experiencing another substitution is only ≈ 1/(1 + 100 × .5) ≈ .02. On the other hand, for the case *p* = .1 we see a modest knee in the time evolution of the fraction of populations that have reverted. This is because a larger fraction ≈ 1/(1 + 100 × .1) ≈ .09 of populations revert before experiencing another substitution, and the subsequent substitution rate is lower, approximately .1 instead of .5.

**Figure 2:**
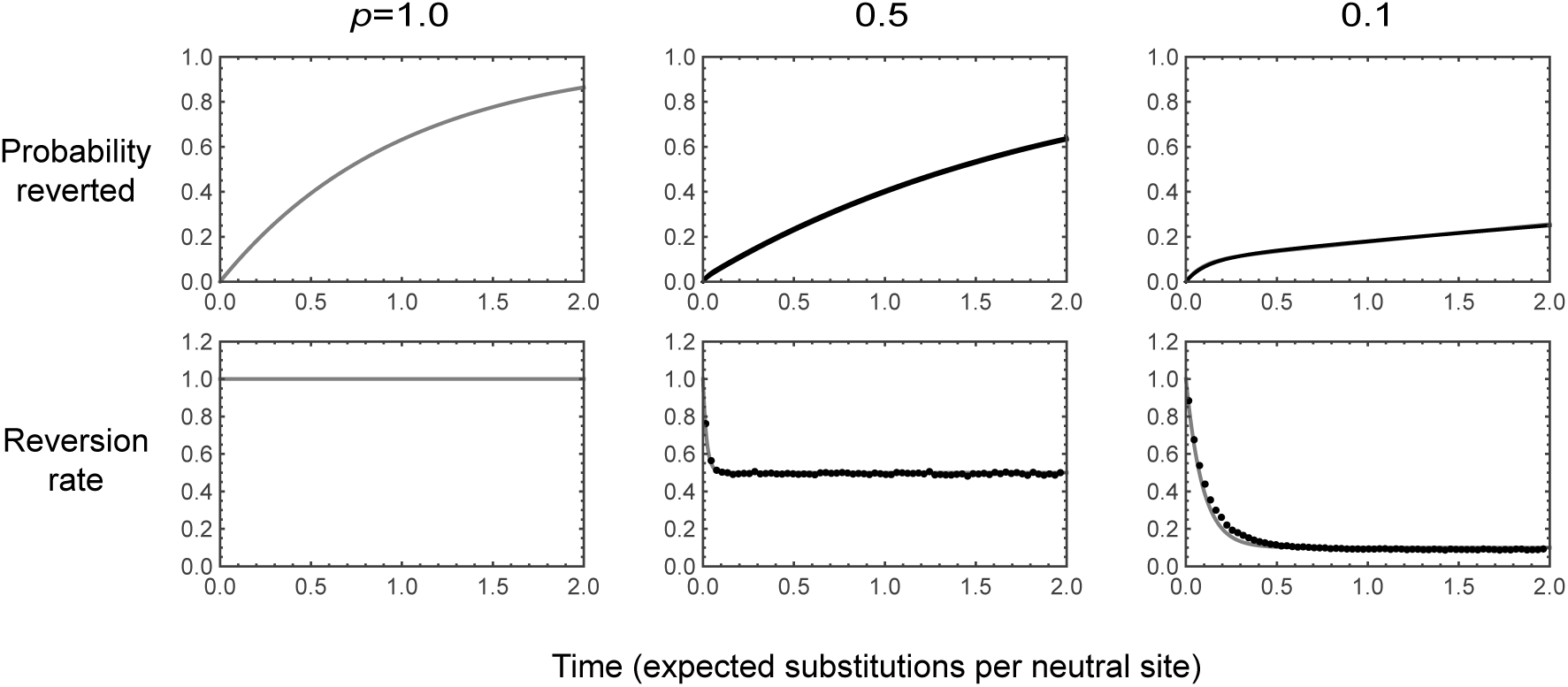
Dynamics of reversion for a biallelic fitness landscape with 100 sites other than the focal site, where each genotype is viable with probability *p*, all viable genotypes are neutral relative to each other, and each site experiences forward and backward mutations at rate 1. The first row shows the probability that a population has reverted as a function of time, and the second row shows the reversion rate. Theoretical results are in gray, and simulations are in black. Dots in bottom row calculate the reversion rate in terms of temporal bins of width .05, where the reversion rate in a bin is calculated as the number of simulations that revert during that bin divided by the average of the number of simulations that have not yet reverted at the beginning of that bin and the number of simulations that have not yet reverted at the end of that bin. Simulation results are based on 1,000,000 runs for each value of *p*.

Intuitively, the knee shape illustrated in Figure 2 arises for small *p* and large *Lp* arises because for small *p* there are fewer other loci at which a substitution can occur, and so populations stay at the genotype they initially arrived at—with its elevated reversion rate—longer, which leads to a substantial probability of having reverted at short times. However, eventually such populations leave the initial genotype, at which time they experience lower reversions rates, ≈ *p*. This produces a pattern where the probability of having reverted becomes substantial at short times but thereafter grows very slowly. Although the theory is complicated and so we do not discuss it in detail here, it is worth noting when *p* is small enough that *Lp* becomes small the waiting time until reversion again becomes short. Intuitively, this occurs because for very small *Lp* the only viable mutation after a substitution at the focal site is the back mutation at the focal site, and thus essentially all reversions occur at the initial elevated rate.

More generally, this simple model suggests that the dynamics of reversion should depend on the degree of mutational robustness even when fitness is allowed to take more than two values. The key insight is that just after a substitution at a focal site, the population is not at a random genotype, but rather one where the back substitution rate is likely to be unusually high. Now, if there is a high degree of mutational robustness, then the neutral substitution rate is high, and the population will leave this special genetic background quickly (c.f. McCandlish 2013), so that when most reversions occur, the substitution rate for the back mutation is much lower than it was initially. When the degree of mutational robustness is very low, by contrast, the population will often stay at the initial genotype until the reversion occurs by a direct back substitution, which occurs at the initial rate. When the degree of mutational robustness is intermediate we see a combination of these two dynamics: a substantial fraction of populations revert via the direct back substitution from the initial genotype, and another substantial fraction revert at the lower rates that are typical of other genetic backgrounds. This intermediate degree of robustness thus produces a characteristic knee in the time-evolution of the probability that the population has reverted.

### Amino acid reversion in codon models

The two examples we have considered so far describe the dynamics of reversion at a single site in a bi-allelic fitness landscape. For our final example we consider the dynamics to a single amino acid under a codon model. Stop codons are treated as inviable, and mutations occur at each site with equal probability to each of the three alternative nucleotides (*J*ukes and Cantor 1969) with a total rate of 1 so that time is measured in the expected number of substitutions per synonymous site (*K*_*s*_). We implement selection by assuming that genotypes with the focal amino acid are neutral relative to each other and have a scaled selective advantage *S* over all other amino acids.

Figure 3 shows the dynamics of reversion under this model for the neutral case, *S* = 0, and also for the case where the focal amino acid has a moderate selective advantage, *S* = 5 (each amino acid corresponds to one line; the fraction reverted is shown in the first row and *h*(*t*) is shown in the second row). The key insight for understanding these dynamics is that, just after a population leaves the set of codons that code for the focal amino acid, it can still mutate back to the focal amino acid. However, after additional substitutions accrue, the population is likely to be at a genotype that is no longer mutationally adjacent to the focal amino acid, leading to a decreasing reversion rate over time. In particular, for both *S* = 0 and *S* = 5, the reversion rate drops from its initial value to nearly its asymptotic value by *K*_s_ = 1. In the case of *S* = 5, this produces a pronounced knee in the shape of the curve describing the time-evolution of the probability that a population has reverted: more than a third of populations are expected to revert almost immediately, wheres the remaining populations take a long time before they eventually revert.

**Figure 3:**
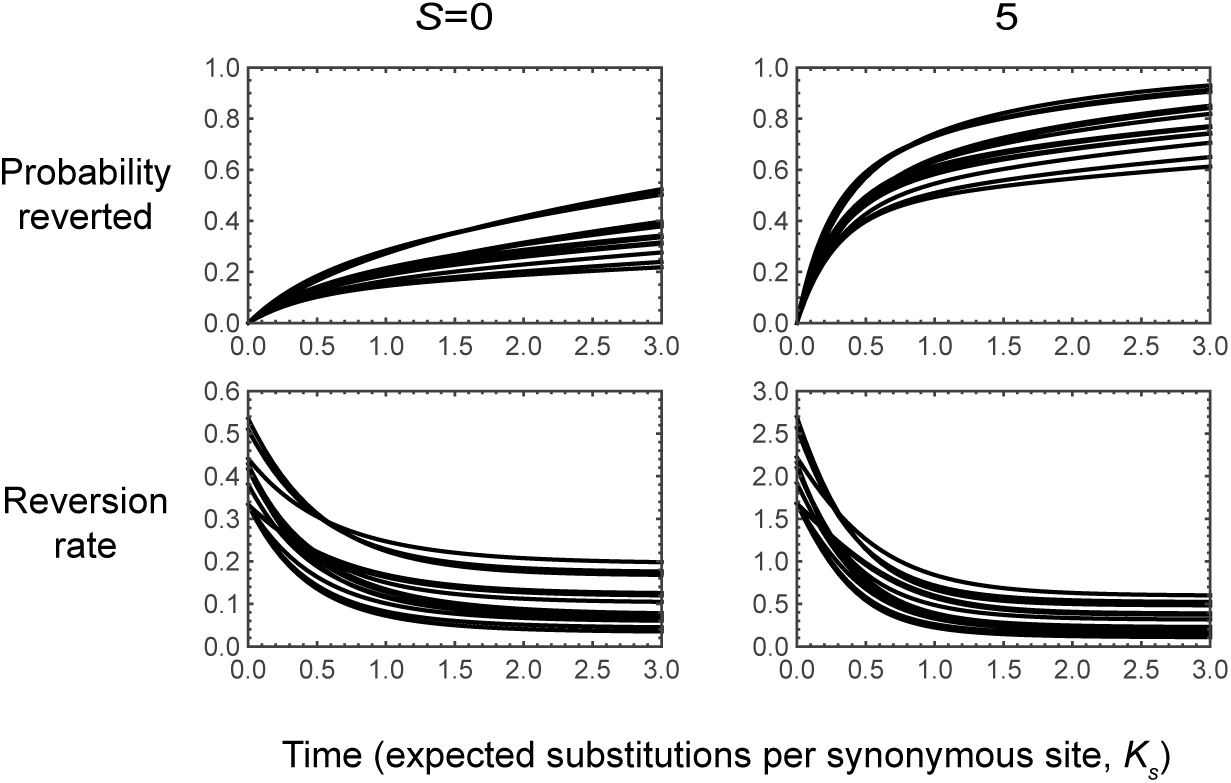
Dynamics of reversion for a codon model where the focal amino acid has scaled selection coefficient *S* over all other amino acids and mutations occur at rate 1 at each site with an equal probability of producing each of the alternative nucleotides. The top row shows the probability of having reverted by time t and the bottom row shows the reversion rate, *h*(*t*), where each of the 20 curves corresponds to a different choice of one of the 20 amino acids as the focal amino acid. Under moderate selection for the focal amino acid (*S* = 5), the time-evolution of the fraction of populations that have already reverted shows a pronounced *k*nee.

## Discussion

We have analyzed reversions in a population that has just left some focal set of genotypes *A*. Our main interest has been the rate at which the population first returns to the set A—that is, the rate at which the population experiences a reversion—and how this rate changes over time. For a population that has already been evolving on a constant fitness landscape for a very long time, we have show that the rate of reversion to *A* is non-increasing in time. Furthermore, when the set *A* consists of all genotypes with a particular allele at a focal site in a bi-allelic fitness landscape, then we showed the reversion rate is strictly decreasing if and only if epistasis is present at the focal site. We have also explored some simple examples of these reversion dynamics, to gain intuition about their magnitude and evolutionary impact.

We have also analyzed the case of a population that has only been evolving on the fitness landscape for a short time, and whose initial state is therefore not drawn from the stationary distribution of the evolutionary dynamics. Here we have shown that the time-evolution of the reversion rate is mathematically identical to the time-evolution of the mean Malthusian fitness for an infinite population evolving on an altered fitness landscape where the substitution rates of the original fitness landscape play the role of mutation rates and the rates of returns to *A* play the role of genotype-specific death rates.

One consequence of reversion rates that decrease in time is a distinctive pattern, whereby a population either reverts very quickly after the initial substitution with some moderate probability or else takes a very long time to revert. If we consider probability of having reverted as function of time, such a pattern can be seen as a “knee”, where this curve rises rapidly at short times until the probability of having reverted is substantial, and then shifts to rising much more slowly. This pattern occurs because the expected rate of substitutions back to a at stationarity is particularly high if we condition on just having left a. As time passes and other substitutions accrue, the features of the genetic background that resulted in the unusually high substitution rate to a are lost. One consequence of this knee-like phenomenon is that the mean waiting time until reversion may not be very informative about the actual dynamics of reversion.

Our results show that at stationarity, a strictly decreasing reversion rate occurs whenever there is genotype-to-genotype variation in the rate of substitutions to the set *A*. When the set a corresponds to the set of genotypes with a particular allele at a particular site, the presence of an epistatic interaction between the focal site and at least one other site is sufficient to produce such genotype-specific variation, and hence sufficient for the reversion rate to be strictly decreasing.

However, other factors besides epistasis can also be responsible for genotype-to-genotype variation in the rate of substitutions to *A*. For instance, in our codon example, which has more than two alleles per site, the structure of the genetic code itself results in variation in the substitution rate to any particular amino acid, because the focal amino acid will not be mutationally accessible from all other codons. More generally, the genotype-phenotype map will tend to produce genotype-to-genotype variation in the rate at which the focal phenotype becomes fixed in the population, so that the reversion rate for phenotypes should typically be decreasing. Finally, it is worth mentioning that while we define a site at the genotypic level by the feature that its mutational dynamics are independent of the states of the other sites, context-dependent mutation could also produce genotype-to-genotype variation in rates of substitution to *A*, which would again be sufficient for decreasing reversion rates (provided the resulting Markov chain is still reversible).

It is also interesting to ask the converse question: what conditions could possibly produce reversion rates that increase in time? To do this, it is helpful to consider a way of restating our main result: if evolution can be modeled as a reversible Markov chain on the set of genotypes, then at stationarity the reversion rate to any subset of genotypes is non-increasing with the time since the population left that subset. A reversible Markov chain is a Markov chain such that the probability of moving through any cycle of states in one direction is equal to the probability of moving around that cycle in the reverse direction (this is known as Kolmogorov’s criterion, see e.g. Kelly 1979, Section 1.5). Thus, a reversible Markov chain is simply a Markov chain with no cyclic biases. This means we can restate our main result as saying that reversion rates can increase in time, starting from stationarity, only if some factor induces a cyclic bias in the Markov chain describing evolution among genotypes.

It is known that, under weak mutation, adding frequency and time-independent natural selection does not add any cyclic bias if no such bias is present in the mutational dynamics (i.e. the weak mutation Markov chain under constant frequency independent selection is reversible if the mutational dynamics are reversible, see e.g. Sella and Hirsh 2005). However, we may expect reversion rates to sometimes increase in time under conditions where fitness is non-transitive (e.g. Kerr et al. 2002) or when environments change in a cyclic manner (e.g. Leslie et al. 2004; Hensley et al. 2009; Bergland et al. 2014). Under these types of conditions, natural selection tends to push populations through a cycle of genotypic states in a periodic manner, which means that reversion becomes more likely as time passes.

Another condition that might produce increasing reversion rates is when the evolutionary dynamics are not stationary, for instance at the beginning of an adaptive transient. We have provided intuition for such circumstances by noting that, under time‐ and frequency-independent selection, the time evolution of the reversion rate can be recast as the time-evolution of the mean fitness of an infinite population on an alternative fitness landscape. This intuition suggests that the reversion rate should tend to go down just as the mean fitness tends to go up, unless the infinite population starts at an unusually high fitness or the reversion rate starts at an unusually low value. In the case of an adaptive transient, early adaptive substitutions will tend to be strongly favored by natural selection meaning that the initial reversion rate is unusually low. In these circumstances we may expect the reversion rate to increase, particularly in the case of diminishing-returns epistasis, where mutations that have strong positive effects early in adaptation have much smaller effects towards the end of adaption (Draghi and Plotkin 2013; Kryazhimskiy et al. 2014, c.f. Hartl et al. 1985; Hartl and Taubes 1996).

Our results help bring clarity to a recent controversy concerning site-specific amino acid preferences during protein evolution (Pollock et al. 2012; Naumenko et al. 2012; Ashenberg et al. 2013; Pollock and Goldstein 2014; Risso et al. 2015; Goldstein et al. 2015; Shah et al. 2015; Usmanova et al. 2015; Bazykin 2015). First, our results show that in the presence of epistasis, reversion rates will be decreasing in time and that the longer a population has left a set of genotypic states, the longer the expected time until reversion. However, our results show that when considering reversion to an amino-acid state, these results already hold even if there is no epistasis at the amino-acid level, because of the structure of the genetic code itself. Indeed, it is worth noting that the epistasis already present in the genetic code is sufficient to produce many of the dynamical signatures of epistatic evolution even if fitnesses are additive at the amino-acid level. For instance, it is easy to specify site-specific amino acid preferences that produce multi-peaked fitness landscapes at individual codons; such codons can have extremely long equilibration times, contrary to the analysis of Kondrashov et al. (2010) and Breen et al. (2013) who only consider the case where every amino acid can mutate to every other amino acid. Overall, decreasing reversion rates are a generic feature of evolution under long-term purifying selection and not a definitive signature of epistasis at the amino-acid level.

Second, it is helpful to distinguish between what is expected when we consider a population that is conditioned to remain outside a subset of states versus one that is simply restricted from entering that subset (c.f. Ashenberg et al. 2013; Pollock and Goldstein 2014). The difference is whether, after a substitution, one only considers populations that have not yet reverted or whether all populations are prevented from reverting (but where we nonetheless keep track of the average propensity to return to the focal subset if such substitutions were to be permitted). We have shown that these two processes have different, but related, mathematical characteristics. If there is any variation in the genotype-specific rates of return to the focal subset then the reversion rate will be strictly decreasing for both processes. While acclimatization of the rest of the genome to being in a new region of the fitness landscape contributes to decreasing reversion rate for both processes, for the conditioned process there is also a statistical effect because populations that spend more time at genotypes with rapid rate of return to the focal subset are likely to revert quickly. In addition, there is a simple mathematical relationship between these two processes: the asymptotic rate of transitions back to the focal subset when populations are restricted from entering it is the rate of an exponential distribution with mean equal to the expected reversion time, and this rate is bounded between the initial reversion rate and the asymptotic reversion rate when conditioning. Thus, while (Ashenberg et al. 2013) have criticized simulations where reversions are prevented from occurring as being unrealistic and misleading, such simulations are in fact more relevant to understanding the evolutionary process than would appear at first glance.

Finally, because of non-linearity in the probability of fixation and genotype-to-genotype variation in mutation rates, the dynamics of the reversion rate and the dynamics of the expected selection coefficient of a reversion mutation may have qualitatively different shapes. Nonetheless, it is the substitution rate that is most closely related to the evolutionary dynamics. This is why we can derive strong results on the reversion rate, but not on the time-evolution of the average selection coefficient of reversions.

## Acknowledgements

This work was funded by the Burroughs Wellcome Fund, the David and Lucile Packard Foundation, US Department of the Interior Grant D12AP00025, and Foundational Questions in Evolutionary Biology Fund Grant RFP-12-16, NIH training Grant 2T32AI055400-11 and US Army Research Office Grant W911NF-12-1-0552.

## Appendix 1 The distribution of reversion times at stationarity

Our analysis of the distribution of reversion times at stationarity relies on the fact that the matrix Q, which givesthe rates for the absorbing Markov chain describing the evolution dynamics until the time of reversion, admits an eigendecomposition. We will develop this eigendecomposition and its properties, and then use these results to show that the distribution of reversion times at stationarity can be expressed as a mixture of exponential distributions. We will then use a very similar analysis to understand the modified process where reversion events are not permitted to occur.

We derive the eigendecomposition of **Q** by first noting some features of the larger matrix **Q**_full_. Because the Markov chain defined by **Q**_full_ is reversible it satisfies the detailed balance relation ***π***_full_(*i*)**Q**_full_(*i*, *j*) = ***π***_full_(*j*)**Q**_full_(*j*, *i*) for all *i*, *j*. As a consequence, the matrix 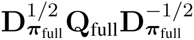 is symmetric, where D_y_ denotes the diagonal matrix with the vector y as its main diagonal.

Now, define the vector ***π*** of length *n* = |*A*^*c*^| such that *π*(*i*) = *π*(*i*)/∑_*j*∈*A*^*c*^_ ***π***_full_(*j*) for *i* = 1,…, *n*. The matrix 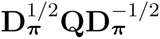 is thus symmetric, since it is simply a constant times a diagonal block of the symmetric matrix 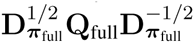. We can thus expand 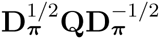 in terms of its eigenvalues and eigenvectors as

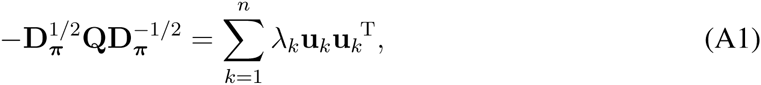

where 0 < *λ*_1_ ≤ *λ*_2_ ≤ *λ*_3_ ≤…< *λ*_*n*_ are the eigenvalues of 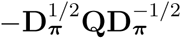 and the eigenvectors **u**_*k*_ are orthonormal (the eigenvalues are real because 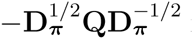 is symmetric and negative because they are same as those of –**Q** where **Q** is the generator of an absorbing Markov chain so that all of its eigenvalues have negative real parts).

We can now use this decomposition of 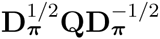 to likewise decompose **Q**. In particular, multiplying Equation A1 by 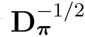 from the left and 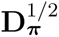 from the right gives us:

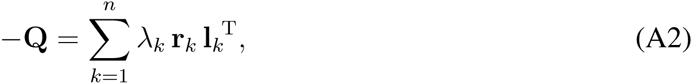

where 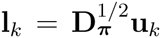 and 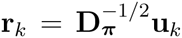 are the left and right eigenvectors of ‒**Q** associated with *λ*_*k*_.

Using this decomposition, we can then write the probability density function of the distribution of reversion times at stationarity as

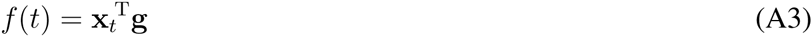

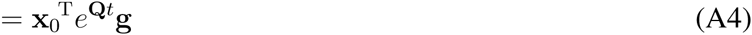

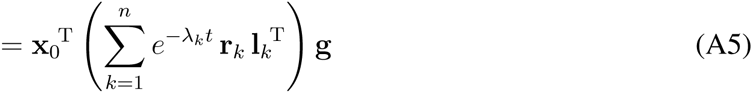

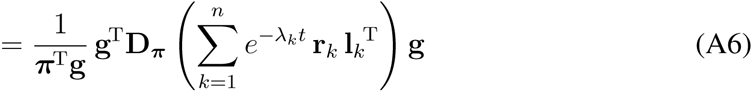

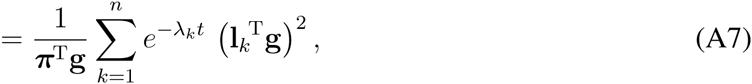

where we have used the fact that at stationarity x_0_(*i*) ∝ ***π***(*i*)**g**(*i*) (Equation 7). Thus, *f*(*t*) is a mixture of exponential densities with rates *λ*_1_,…, *λ*_*n*_ where the exponential density with rate *λ*_*k*_ has weight (**l**_*k*_^T^g)^2^/(*λ*_*k*_ **π**^T^g).

We just showed that the distribution of reversion times at stationarity is a finite mixture of exponential distributions. We will now show that the hazard function for any finite mixture of exponential distributions is non-increasing. In particular, if *f*(*t*) is a finite mixture of exponen-tial densities

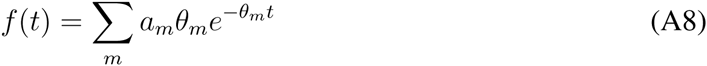

with *a*_*m*_, *θ*_*m*_ > 0, *∑*_*m*_ *a*_*m*_ = 1, then the hazard function is

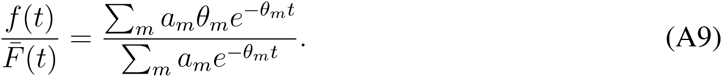

Differentiating the hazard function, we see that that the sign of the derivative depends only on the sign of

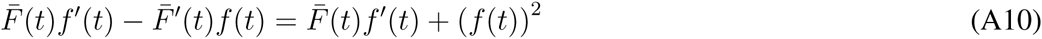

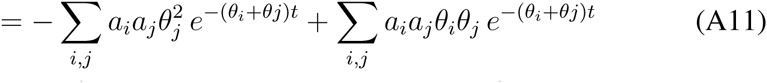

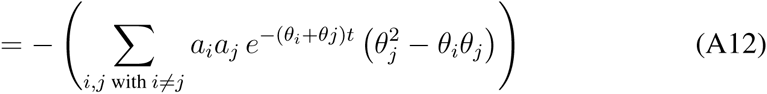

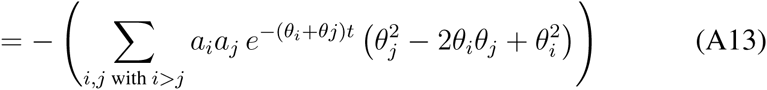

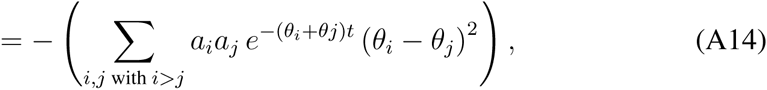

which is non-positive since each term in the sum is a product of non-negative quantities and hence non-negative. Thus, the hazard function is non-increasing.

We turn now to the analysis of the modified process with rate matrix **Q**\* = **Q** + **D**_g_, where **D**_g_ is the diagonal matrix with g on its main diaogonal. Let us start by assuming that *A*^*c*^ is mutationally connected; we will return to the case of disconnected *A*^*c*^ momentarily. First note that **Q**\* only differs from **Q** on its diagonal entries. Thus, following our analysis for **Q**, 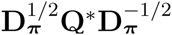 is symmetric and can be expanded in terms of its eigenvalues and eigenvectors as

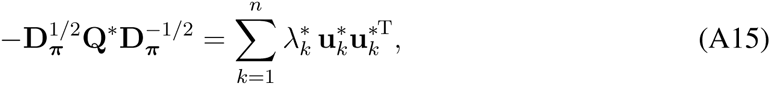

where 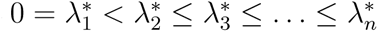 are the eigenvalues of 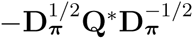 and the eigenvectors **u**^*k*^ are orthonormal (we have 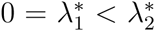 because *A*^*c*^ is connected and so the continuous time Markov chain generated by **Q**\* is ergodic). With this decomposition in hand, we can write 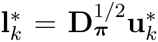 and 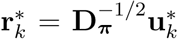 as the left and right eigenvectors of –**Q**\* associated with 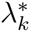 and write the reversion rate under the modified process as:

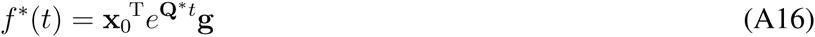

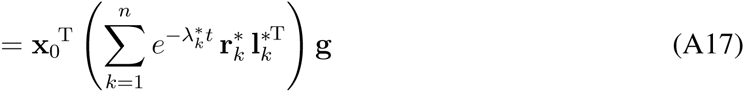

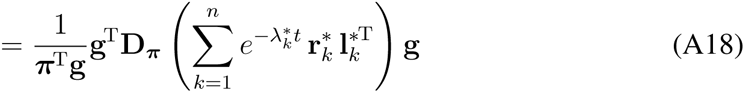

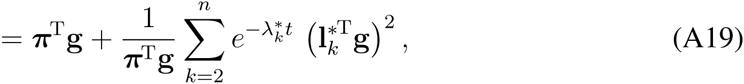

where we’ve used the fact that the row sums of **Q**\* are all 0 so that 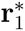 is the vector of all 1s and thus 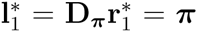. Because 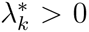 for *k* ≥ 2, this expression is clearly strictly decreasing in t unless *n* = 1 (*i*n which case **g**(*i*) is obviously constant) or 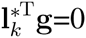 for *k* ≥ 2. In the latter case, we have 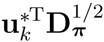 for all *k* ≥ 2, so that 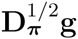 must be orthogonal to all the 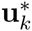 for *k* ≥ 2. Since the 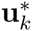 are an orthonormal basis of ℝ^*n*^, the only direction remaining for 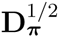 is 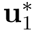, so that g must be in the direction of 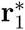, which is the vector of all 1s. Thus, in that case g is also constant.

So far, we’ve shown that provided that *A*^*c*^ is mutationally connected, at stationarity the reversion rate under **Q**\* is constant if **g**(*i*) is constant and otherwise it is strictly decreasing. Now we consider the case where *A*^*c*^ has several disconnected components, *A*_1_,…*A*_*i*_. Clearly the total reversion rate is a weighted average of the reversion rates for each disconnected component when analyzed separately as above. If **g**(*i*) is non-constant within any one of these components, than that term in the average is strictly decreasing in time and hence the reversion rate as a whole is strictly decrease. Thus, for general *A*^*c*^ the reversion rate at stationarity under **Q**\* is constant if **g**(*i*) is piece-wise constant in the sense of being constant for each of the disconnected components of *A*^*c*^, otherwise it is strictly decreasing.

## Appendix 2 Role of epistasis in multiallelic fitness landscapes

In the main text, we showed that for biallelic fitness landscapes, the reversion rate at stationarity for a site is a decreasing function of time if and only if that site is involved in an epistatic interaction with another site. Here we show that for multi-allelic fitness landscapes the presence of an epistatic interaction with another site is a sufficient condition under stationarity for reversion rates at the focal site to be decreasing.

First we must establish some notation for multi-allelic fitness landscapes. If we have a fitness landscape over *L* sites, we can label the *n*_*l*_ ≥ 2 alleles at the *l*-th site, *α*_*l*,1_,…, *α*_*l*,*n*_*l*__ and write the mutation rate between *α*_*l*,*k*_ and a_*l*,*m*_ as *μ*_*α*_*l*,*k*__→*α*_*l*,*m*_. Thus, if genotypes *i* and *j* differ at more than one site we have *M*_full_(*i*, *j*) = 0 otherwise they differ at exactly one site / in which case *M*_full_(*i*, *j*) = *μ*_*α*_*i*,*k*_→*α*_*l*,*m*__ where genotype *i* has the *α*_*l*,*k*_ allele at site *l* and genotype *j* has the *α*_*l*,*m*_ allele at site *l*.

Without loss of generality we can define the focal set a to be the set of genotypes with allele *α*_1,1_ in site 1. We want to show that if site 1 has an epistatic interaction with at least one other site, then the **g**(*i*) are not constant, since this is sufficient to show that the reversion rate is decreasing at stationarity.

Now, either *μ*_*α*_1,*k*_→*α*_1,1__ > 0 for all *k* or not. If not, then *μ*_*α*_1,*k*_→*α*_1,1__ > 0 for some *k*, since otherwise the Markov chain defined by Qfull would not be reversible (populations could never return from *A*^*c*^ to *A*). Now, if *μ*_*α*_1,*k*_→*α*_1,1__ > 0 for some *k* but *μ*_*α*_1,*m*_→*α*_1,1__ = 0 for some *m*, then g is positive for genotypes with allele *α*_1,*k*_ at site *l*, but 0 for genotypes with allele *α*_1,*m*_ at site *l*. Thus, g is non-constant.

We thus turn to the case where *μ*_*α*_1,*k*_→*α*_1,1__ > 0 for all *k*. Our strategy is to show that if all alleles at a site can mutate to all the other alleles, then the presence of any epistatic interaction between one site and another is sufficient, by the transitivity of fitness differences, to guarantee the existence of a set of four genotypes to which we can apply the argument in the main text for biallelic sites. Now, if site 1 is epistatic with some other site, then without loss of generality we can let this other site be site 2. Because site 1 is epistatic with site 2 there exist, by the definition of epistasis, genotypes *i*, *i*′, *j*, *j*′ and alleles *α*_1,*k*_, *α*_1,*m*_, *α*_2,*k*′_, *α*_2,*m*′_ such that:

1. Genotype *j* and *i*′ are both single mutants with respect to genotype *i*, and *j*′ is the corresponding double mutant. In particular, genotype *j* is formed from genotype *i* by a single *α*_1,*k*′_ → *α*_1,*m*_ mutation, genotype *i* is formed from *i* as a single *α*_2,*k*_ → *α*_2,*m*_, and the double mutant genotype *j*′ is formed from genotype *i* by the combination of an *α*_1,*k*_ → *α*_1,*m*_ mutation and an *α*_2,*k*_ → *α*_2,*m*_ mutation.
2. **f**(*i*) – **f**(*j*) ≠ **f**(*i*′) – **f**(*j*′).

If either *k* = 1 or *m* = 1 then g is non-constant by the same argument as given for the two allele case in the main text. Therefore suppose *k* = 1 and *m* = 1. In that case we can find genotypes *h* and *h*′ such that *h* is identical to *i* and *j* except that it has the α_1,1_ allele at site 1 and likewise *h*′ is identical to *i*′ and *j*′ except that it has the α_1,1_ allele at site 1. Now, if it were the case that both **f**(*h*) – **f**(*j*) = **f**(*h*′) – **f**(*j*′) and **f**(*i*) – **f**(*h*) = **f**(*i*′) – **f**(*h*′), then we would have

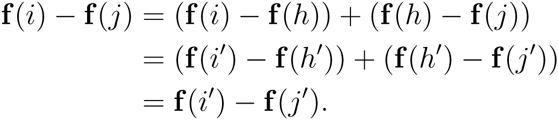

But **f**(*i*) – **f**(*j*) = **f**(*i*′) – **f**(*j*′), so either **f**(*h*) – **f**(*j*) = **f**(*h*′) – **f**(*j*′) or **f**(*i*) – **f**(*h*) = **f**(*i*′) – **f**(*h*′). Thus, we could have chosen *h* and *h*′ instead of either *i* and *i*′ or *j* and *j* above, in which case the non-constancy of g again follows by the same argument given for the two allele case in the main text.

## Appendix 3 Expected time until reversion

We want to derive the expected time until reversion to a set *A* at stationarity given that the population has just left *A*. To study this time, it is helpful to define a modified Markov chain on a state space *A*^*c*^ ∪ {*A*} where the new state *a* takes the place of the set *A* in the original chain.

In particular, consider the chain with rate matrix

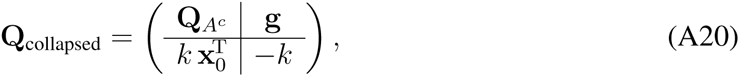

where *k* is the expected substitution rate from a to *A*^*c*^ at stationarity for **Q**_full_ conditional on the population being in *A*, and x_0_(*i*) is the probability of the population being at genotype *i* ∈ *A*^*c*^ under **Q**_full_ at stationarity, conditional on a substitution from *A* to *A*^*c*^ having just occurred. By construction, this new chain with rate matrix **Q**_collapsed_ has the same distribution of reversion times to *a* at stationarity as did the original chain.

Furthermore, if *A*^*c*^ contains n elements, it is easy to verify that the stationary distribution of **Q**_collapsed_ is given by the vector (***π***_full_(1),…, ***π***_full_(*n*), ***π***_full_(*A*)), where ***π***_full_(*A*) = ∑_*i*∈*A*_*π*_full_(*i*), since the new chain conserves both the stationary rate of substitutions from *a* to *i* and from *i* to *A* for all states *i* ∈ *A*^*c*^.

Analyzing this new chain, a straightforward application of the reward-renewal theorem for the renewal process defined by returns to a and reward equal to the waiting time before leaving *a* after each return gives us:

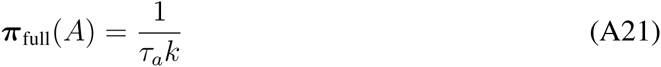

(Grimmett and Stirzaker 2001, Theorem 10.5.10), where *τ*_*a*_ is the expected waiting time for a population currently at *a* to leave a and then return to *a* for the first time.

Now, a population leaving a and then returning to a must first leave a and then revert to a. The expected time before the population leaves a is 1 /*k* and then the expected waiting time to revert to a is *m*(0), the quantity that we are trying to develop an expression for. Thus, we have

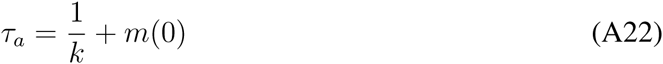

Plugging Equation A22 into Equation A21 and solving for *m*(0) gives us:

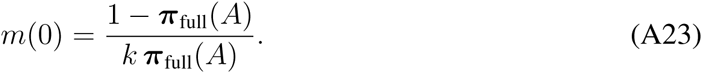

At stationarity, the total rate that populations arrive at a must be equal to the rate that they leave a, so we have *k* ***π***_full_(*A*) = ∑,_*i*∈*A*_*c* ***π***^full^(*i*)**g**(*i*), and of course 1 – ***π***_full_(*A*) = ∑_*i*∈*A*^*c*^_***π***(*i*).

Substituting these expressions into Equation A23 yields

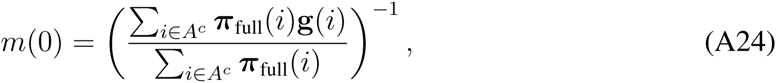

i.e. the expected waiting time reversion is equal to the reciprocal of the expected substitution rate to *a* at stationarity conditional on the population being in *A*^*c*^, as required.

